# Evidence for grid-cell-related activity in the time domain

**DOI:** 10.1101/2022.06.14.476894

**Authors:** Gregory Peters-Founshtein, Amnon Dafni-Merom, Rotem Monsa, Shahar Arzy

## Abstract

The relation between the processing of space and time in the brain has been an enduring cross-disciplinary question. Grid cells have been recognized as a hallmark of the mammalian navigation system, with recent studies attesting to their involvement in organization of conceptual knowledge in humans. To determine whether grid-cell-like representations support temporal processing, we asked subjects to mentally simulate changes in age and time-of-day, each constituting “trajectory” in an age-day space, while undergoing fMRI. We found that grid-cell-like representations supported trajecting across this age-day space. Furthermore, brain regions concurrently coding past-to-future orientation positively modulated the magnitude of grid-cell-like representation in the left entorhinal cortex. Our findings suggest that temporal processing may be supported by spatially modulated systems, and that innate regularities of abstract domains may interface and alter grid-cell-like representations, similarly to spatial geometry.

## Introduction

The human experience unfolds in space and time. Several lines of research have suggested that these two domains are intimately related within the neurocognitive system (1, 2). For instance, the representation of time is subjected to spatial priors, as evident in everyday language (e.g. “I look *forward* to meeting you”; 3–5). The fundamental human capacity to “mentally time travel” (6, 7), i.e recalling past- and simulating future events, is underlain by categories of space and time (8). Moreover, mental time travel has been proposed to “spatialize” along a “mental timeline”(1,9), akin to the mental number line (10). Correspondingly, cognitive processes involving the mental time line have been shown to be influenced by spatial biases, such as the direction of writing (11, 12) or manipulations of spatial attention (13, 14).

The discoveries of place and grid cells have provided a physiological basis to the representation and association between space and time, bridging neurocognitive models and neurophysiological correlates (15–17). Place cells, neurons which code for specific places in the environment, were first reported in the rodent (18) and subsequently in the human hippocampus (19). In recent years, studies have shown that neurons in the rodent (20) and human (21) medial temporal lobe (MTL) also have robust temporal correlates. Analogously to the association between place cells’ firing patterns and specific locations, “time cells” were found to reliably fire at specific and consistent moments within a longer time interval (20). Moreover, a direct comparison between different task conditions revealed neurons which interchangeably code for time or place (22, 23). Years after the discovery of place cells, grid cells were recorded from the entorhinal cortex (EC) and named for their hexagonally symmetric firing patterns (16, 24, 25). Unlike place cells, grid cells are thought to retain their regular spatial structure, scale, and position relative to other grid cells across environments and as such are thought to code the metrics of the spatial cognitive map (26, 27).

In recent years, researchers have used virtual navigation tasks to show that hexagonally symmetric grid-cell-like representations could be measured by functional MRI (fMRI) (28, 29). Ensuing studies have demonstrated that such grid-cell-like representations support not only spatial navigation, but also abstract cognitive processes (30). Additional studies showed grid-cell-like representations when “navigating” two-dimensional visual (31, 32), auditory (33), olfactory (34), social hierarchy (35) and semantic spaces (36). However, it remains to be determined whether grid-cell-like representations may also support temporal processing (37).

In this study, we sought to determine whether spatially modulated mechanisms contribute to humans’ ability to mentally traject across time. We used fMRI to establish whether humans evoke grid-cells-like, hexagonally symmetric, representations when imagining gradual changes in time. For this purpose, we designed a task analogous to that used for navigation in physical space (38), with the dimensions of age and time-of-day rather than physical space. We further inquired whether the past-to-future orientation of the mental timeline could interface with the hexagonally symmetric grid-associated signal.

## Results

### Trajections across a day-age space evoke grid-cell-like representations in the EC

To examine whether grid-cell-like representations in the EC support the ability to mentally traject across time, 28 healthy subjects gained proficiency in “navigating” a two-dimensional space, spanned by changes in age and time-of-day (“age-day space”, Fig. 1). Each point on this age-day space represents a unique intersection of age and time-of-day. Specific age-day points are associated with related symbols, representing characteristic events (“event symbols”). For example, a child cycling in the afternoon or an older person using binoculars while attending an evening show at the theater (Fig. 1). Following four days of training (for training protocol and performance see supplementary methods and results), subjects performed an fMRI-adapted task (Fig. 2), in which they watched videos depicting changes in age and time-of-day simultaneously, each according to a predefined age:day ratio (Fig. 2). Each video corresponded to one “trajectory” in the age-day space and consisted of several stages: (1) morphing, (2) imagination and (3) feedback (Fig. 2A,B, Movie S1). During “morphing”, subjects viewed a single age-day trajectory for 8 seconds. Next, during “imagination”, subjects mentally simulated morphing at the same age:day ratio for an additional 4 seconds. Finally, during “feedback”, subjects were presented with three different event symbols and were instructed to select the one associated with the trajectory’s outcome.

**Figure 1.**
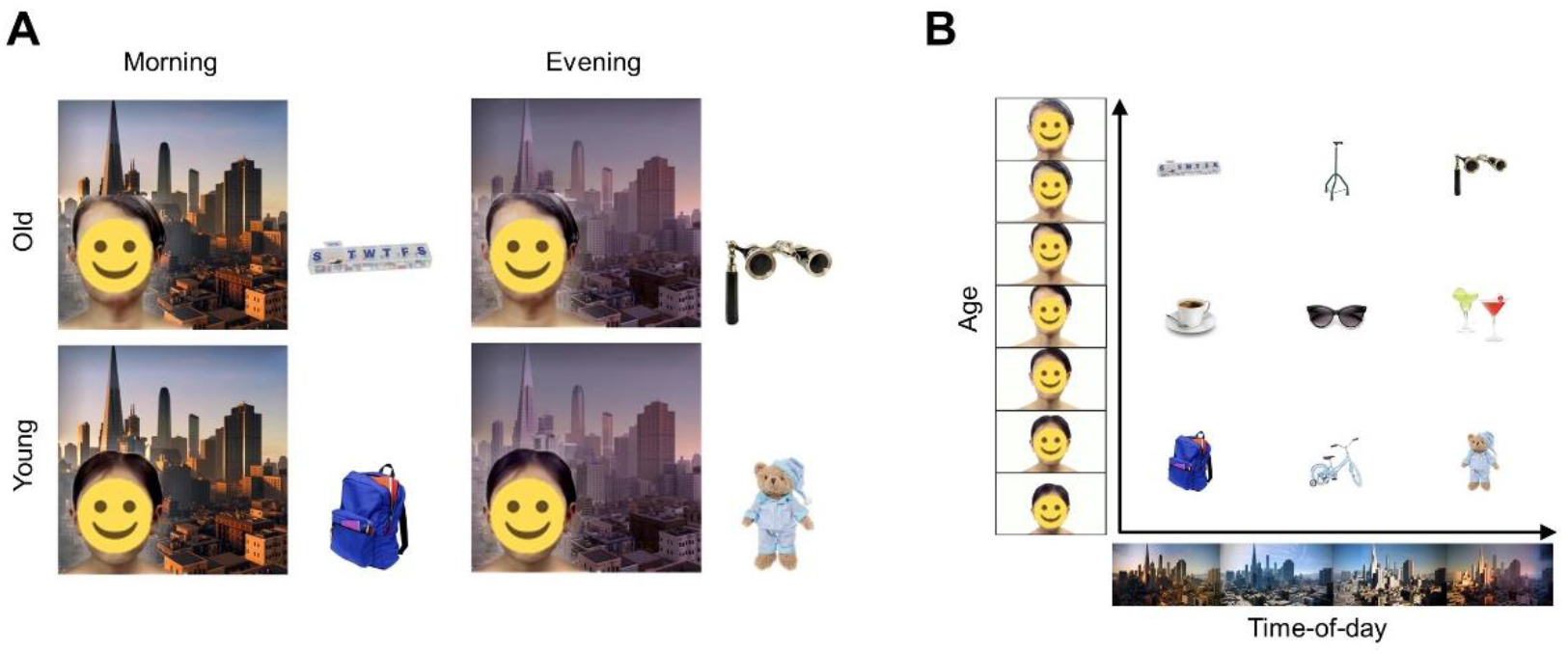
Navigation in a two-dimensional temporal space. (A) Association between event symbols and age-day space. Subjects were trained to associate coordinates in the age-day space (image of person against city backdrop) with event symbols: 9 everyday objects, related to their corresponding age-day coordinates. 4 exemplary pairs of event symbols and age-day coordinates are presented in a 2×2 matrix. **(B)** Age-day space with embedded event symbols. Visual representation of the “two-dimensional temporal space”, indicating the range of variability across the age variable (y axis) and the time-of-day variable (x axis), as well as the 9 embedded event symbols. Importantly, subjects were not consciously aware that these associations could be organized in a continuous two-dimensional space.

**Figure 2.**
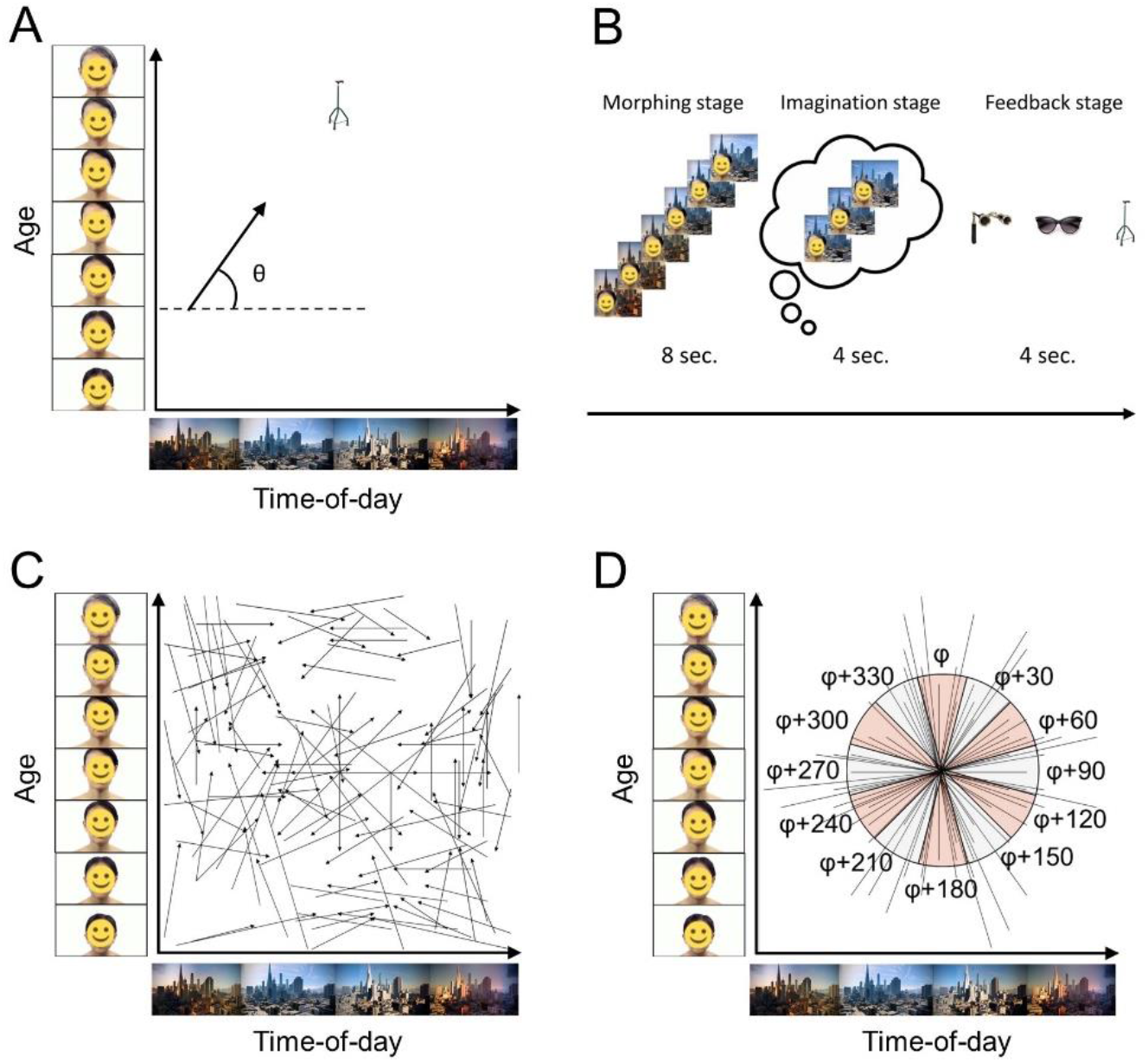
Experimental design for navigation in the time domain. **(A)** Example of a trajectory in the age-day space. A “location” in this abstract space was represented by a specific age and time-of-day combination. A trajectory was equivalent to visually continuously changing the age and time-of-day. The direction θ of the trajectory depended on the ratio of the rates of change of age and the time-of-day (Movie S1). **(B)** An exemplary trial, corresponding to the trajectory in panel A and consisting of 3 stages: (1) visual “morphing”; (2) “imagination” (mentally morphing) and (3) “feedback” (selecting the appropriate outcome event symbol). **(C)** Layout of all trajectories across the age-day twodimensional space. **(D)** Visualization of “aligned-misaligned” analysis. Visualizing a core principle of the analysis, the origins of all trajectories were shifted to a single point while maintaining their original orientations. These trajectories were further categorized as either aligned (orange sectors) or misaligned (grey sectors) with relation to the mean orientation angle ϕ of the hexagonal grid. Note that ϕ is different for each participant (see methods section for details on how ϕ was calculated). Here, for example, the direction θ is misaligned with relation to the grid.

Performance was assessed during the “feedback” stage. Success rates (Mean±standard error of the mean (SEM), 0.68±0.02) were significantly higher than chance (0.33, one-sample t-test, t(27)= 21, p<0.001). This indicates that subjects have established a representation of the age-day space and the embedded event symbols.

To test whether trajections in time evoked hexagonally symmetric grid-cell-like representations in the EC, we first identified EC voxels with maximal hexagonal modulation (for elaboration see “hexagonal modulation analysis” in Methods). We then used part of our data (run 1) constructed regions of interest (ROIs) around maximally-modulated voxels, and used them in an independent (runs 2,3) split-half analysis, where grid orientation was estimated on half the data and applied to the other half (30). Importantly, to avoid circular inference, localizer and split-half analyses were conducted using different parts of the data. Within the EC ROIs and, split-half analysis showed parameter estimates of aligned trials (Mean±SEM, LH: 0.16±0.4; RH: 0.45±0.53; Fig. 3B) to be significantly higher than those of misaligned trials (Mean±SEM, LH: −0.89±0.6; RH: −0.83±0.64; Fig. 3B) in the EC bilaterally (paired t-test, left: t(27) = 2.14, p<0.05; right: t(27) = 2.21, p<0.05, FDR-corrected; Fig. 3B). Significant differences were found for the 6-fold, hexagonal, symmetry model but not for biologically implausible control models (5-fold, 7-fold, paired t-test, p>0.05; Fig. 3B). These finding show that mentally trajecting across the age-day space evokes EC-based grid-cell-like presentation.

**Figure 3.**
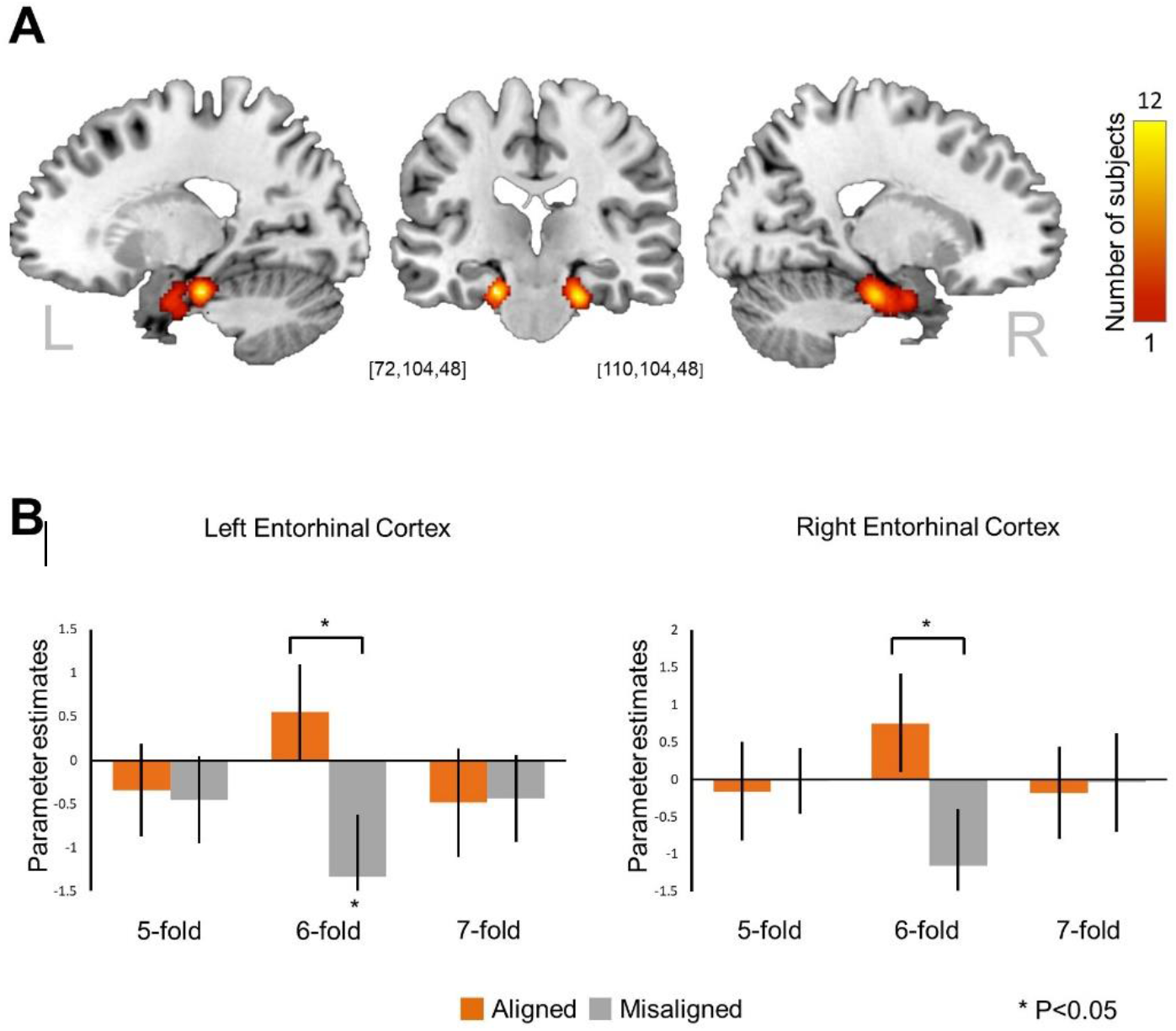
Grid-cell-like representations in the entorhinal cortex (EC). **(A)** Regions of interest for grid-cell-like effect analysis. The EC was segmented individually for every subject. For each subject, within their EC mask, the maximal amplitude (of hexagonal modulation) voxel was calculated from the first run data only and selected as the center of a 6mm-radius sphere. All spheres were jointly superimposed on an MNI brain template and color coded to indicate the degree of overlay between subjects across voxels. **(B)** Estimating the grid-cell-like effect. Using only data from runs 2- and 3 we conducted a split-half analysis, where grid orientation was estimated on half the data and applied to the other half. Split-half analysis showed that parameter estimates of aligned (orange) trials were significantly larger than those of misaligned (grey) trials bilaterally (left EC - t(27) = 1.89, p<0.05; right EC - t(27) = 1.9, p<0.05; FDR-corrected). This pattern was found for 6-fold rotational symmetry but not for biologically implausible control models (5-fold, 7-fold). * P < 0.05. Error bars represent SEM.

### Interactions between grid hexagonality and timeline-congruency

Considering past-to-future orientation of the mental timeline as a core property of human temporal processing, we set to evaluate past-to-future congruency effects on behavioral performance, neural activity and finally grid-cell-like representations. Trials were considered as “congruent” if age-day trajectories maintained past-to-future orientation in both age and time-of-day dimensions, and “incongruent” otherwise. Congruent trials were associated with superior performance, manifesting as both significantly higher success rate (paired t-test, t(27) = 4.83, p<0.001) and significantly lower response times (paired t-test, t(27) = 2.4, p < 0.05), as compared to non-congruent trials (Fig. 4C). To determine if brain activity codes past-to-future orientation, independently of assumptions of hexagonal symmetry, we used a searchlight-based multi-voxel pattern analysis (MVPA, 39). MVPA revealed that multiple brain regions significantly code congruency (Z-score>1.645, P<0.05) including the hippocampus and parahippocampal gyrus bilaterally, as well as right EC, right insula, right operculum and left superior temporal gyrus (Fig. 4D). Importantly, MVPA-generated congruency maps overlapped with the right more than left EC ROIs (Fig. 4E; central voxels: right: 7%; left: 0.4%, whole spheroids: right: 54%; left: 16%).

**Figure 4.**
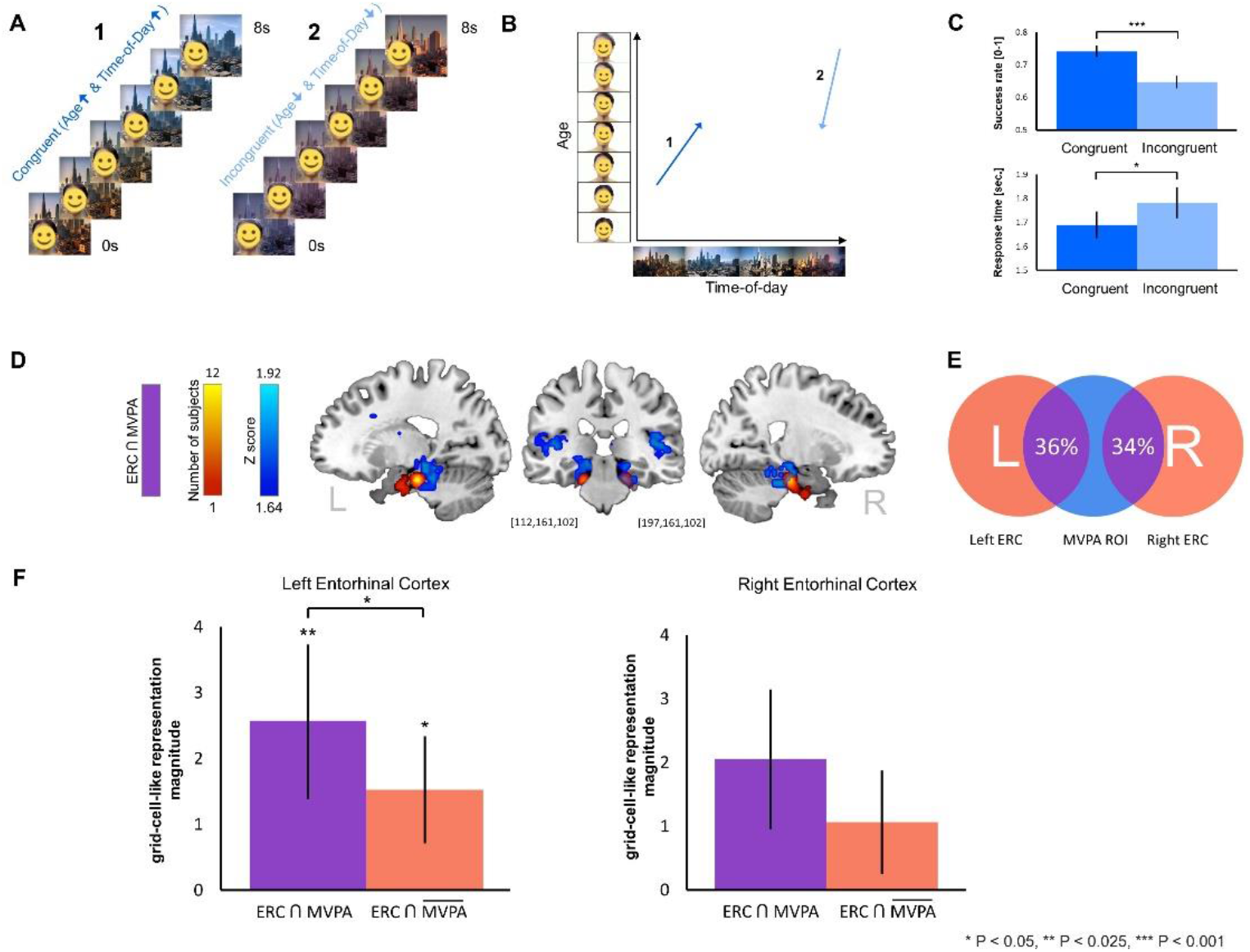
Interactions between timeline-congruency and grid code hexagonality. **(A)** Exemplary pair of past-to-future congruent and incongruent trials. Panels A1 and A2 illustrate the morphing stage of a congruent (A1; young-to-old and morning-to-evening) and incongruent (A2; old-to-young and/or evening-to-morning) trials. **(B)** Exemplary congruent and incongruent trajectories. Exemplary pair of congruent (1, dark blue) and incongruent (2, light blue) trajectories in the age-day space corresponding to the trials in panel A. **(C)** Behavioral effects of future-to-past congruency. Success rates and response times of congruent and incongruent trials were compared in a pair-wise manner to reveal significant benefits in performance for congruent trials in both success rate (p<0.001) and response time (P<0.05). **(D)** Multi-voxel pattern analysis (MVPA) of timeline-congruency. Sagittal (left and right) and coronal (middle) views are overlaid with both EC ROIs (orange), the results of group-level MVPA, decoding trial timeline-congruency (blue) (p<0.001, TFCE-corrected; cluster size threshold: 20 voxels). MVPA – EC overlap regions colored in solid purple. Purple delineation marks all spheroid voxels used in MVPA analyses. **(E)** MVPA and EC overlap. A Venn diagram indicating overlap (purple) between right (34%) and left (36%) of EC ROIs voxel (orange) with MVPA-generated congruency map (blue) on a single subject level. **(F)** Congruency – Hexagonality interaction. Split half grid codes analysis was repeated separately for EC ROIs voxels yielding MVPA probability >0.5 (purple, “EC ∩ MVPA”) and ≤0.5 (orange, 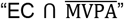), to reveal that the magnitude of grid-cell-like representation is significantly increased in left EC ∩ MVPA over EC ∩ 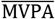 voxels (t(27) = 2.03, p<0.05). * P < 0.05, ** P < 0.025, *** P < 0.001. Error bars represent SEM.

Having shown that past-to-future congruency affects performance, and is coded in brain activity, we inquired whether congruency could modulate grid-cell-like representations. As a preliminary step, we separated EC ROI voxels into “coding” and “non-coding” ROIs, according to their MVPA-derived probability to detect congruency above chance level. Using data from runs 2,3, we then repeated split-half analysis with grid orientation estimated on the first half of the data and applied it to the second half, within congruency coding (EC ∩ MVPA) and non-coding (EC ∩ MVPA) voxels. This procedure, revealed the magnitude of grid-cell-like representations to be significantly increased for congruency-coding over non-coding voxels in the left EC (paired t-test, left EC (t(27) = 4.60, p < 0.025, FDR-corrected; Fig. 4F). Taken together, our findings demonstrate that entorhinal-based grid-cell-like representation can support humans’ ability to mentally traject across time. Moreover, past-to-future congruency affected task performance, was coded in task-evoked activity, and modulated the grid-cell-like representation.

## Discussion

The present work showed humans to form a two-dimensional space as they modified the age and time-of-day parameters of a conjugated stimuli. Trajecting the age-day space while undergoing fMRI, revealed participants to evoke grid-cell-like, hexagonally symmetric, representations. In-depth analysis of the grid-cell-like representations indicated that trajections congruent with past-to-future orientation positively modulated the grid-associated signal within regions coding for congruency. These findings are discussed with respect to the literature concerning cognitive maps and the coding of space, time and abstract concepts.

Time is often regarded as the “4th dimension” to the three spatial ones. The discovery of “time cells”, and more specifically, the potential of neurons to interchangeably code space or time under different experimental conditions (22, 40, 41), may suggest a distribution of labor within the hippocampal-entorhinal system when coding spatiotemporal context (42, 43). Ensuing works extended the scope of domains represented by the grid system to include: gaze movement across the field of view (31, 32), sound frequencies (33), a two-dimensional “odor space” comprised of olfactory stimuli (34), abstract spaces, spanned by the physical properties of bird images (30), and the size and accompanying sound pitch of “audiovisual objects” (36). Considering the diversity of domains comprising this body of work, it is plausible to suggest that the entorhinal grid system provides a domain-general mechanism to code for salient, task-relevant dimensions of experience (17, 44), including time.

How the neurocognitive system marks and represents the flow of time, that is differentiating recalled past events from simulated future scenarios, is a matter of debate. This question grows more difficult when considering the body of evidence recognizing the same neurocognitive system as underlying both recalling the past and simulating the future (45, 46). Several theories attribute this distinction to inherent properties of past and future representations. One theory claims that imagined future events are more emotionally positive than past ones (47–49). Another theory contends that imagining future experiences, but not remembering past experiences, requires flexible reconstruction of recalled details into novel events (50). Finally, individuals contemplate and value the future more than the past (51–53), as the future is determined by their present decisions and is highly connected to their goals (54). Our results, showing that coding past-to-future congruency can significantly modulate the magnitude of grid-cell-like representation, may add to these neurocognitive models. We propose that past-to-future congruency can be “imprinted” onto the grid-code, and thus may generate an inherent preference for past-to-future directed mentations. Further research is needed to examine this hypothesis.

Grid cells are often suggested to provide an “absolute metric” onto which self- and object locations are updated (26, 27). Recent works revealed how geometrical environmental features can influence and distort the rigid grid structure, consequently challenging the characterization of an invariant metric (55, 56). In humans, grid-cell-like representations were found to be modulated by proximity to boundaries (57) and bodily self-consciousness (58). Considering both the potential for signal modulation and domain-generality as core properties of the grid system, we asked whether the grid system retains its specific firing patterns across cognitive spaces. Our results, showing congruency-driven modulation, suggest that as the grid system maps abstract cognitive spaces, it might be distorted through interaction with the “native” abstract geometry, similarly to physical geometry. Corresponding findings demonstrate that nonspatial reward information distorts grid fields, warping them toward goal locations and leading to long-lasting deformations of the entorhinal map (59). As we go forward exploring the contributions of spatially-modulated systems to higher-order cognitive processes, we need to address not only how complex experiences are mapped and coded, but also how their innate regularities might themselves interface with the code.

This study is not free of limitations. As we consider visual information to play an essential role in our perception and processing of time, we’ve designed an ecological visual representation of time and used images representative of their age-day coordinates. However, the visual cues themselves do not inherently represent any episodic information but merely allude to the underlying temporal content. Therefore, generalization beyond those aspects of mental simulation should be made with caution. Additionally, while we designed a user interface devoid of spatial visualizations and avoided spatially related language during the experiment and preceding training, we nevertheless used a sensorimotor device (computer mouse) during training. The sensorimotor spatial input during training might have contributed to the recruitment of the spatially-modulated grid system during the fMRI-adapted task, even as the latter had no visual spatial properties.

In conclusion, this study has demonstrated that grid-cell-like representations were evoked as subjects mentally trajected across a feature space spanned by age and time-of-day, suggesting that the representation of time potentially relies on spatially modulated systems. Past-to-future orientation constrained and modulated task performance and the evoked hexagonally symmetric grid-associated signal. We ascribe this effect to the fact that age and time-of-day are salient features of temporal organization. Finally, considering the grid-system’s role in higher cognitive processes, we propose to explore not only how complex experiences are coded by spatially-modulated systems, but also how the innate regularities of these abstract domains might themselves interface with the code.

## Materials and Methods

### Subjects

28 healthy subjects (15 males, mean age: 26.64 ± 0.57, all right-handed) participated in the study. All subjects reported to be in good health with no history of neurological or psychiatric disease and with normal or corrected-to-normal eye vision participated. All subjects provided written informed consent. The study was approved by the ethics committee of the Hadassah Medical Center.

### Behavioral Training

Subjects performed a series of stimulus-outcome learning tasks. The tasks were all performed using a specially designed interactive image of a person set against a city backdrop (Fig. 1B), in which the person’s age and the backdrop’s time-of-day could be manipulated using mouse movements. Through a series of 4 tasks, subjects learned to readily adjust the age and time-of-day parameters to generate age-day combinations as well as to associate specific age-day combinations with 9 different embedded event symbols (Fig. 1), related to their age-day coordinates. Importantly, while each event image could be effectively described as an embedded stimulus within a two-dimensional environment, subjects were not consciously aware of the underlying two-dimensional structure. Images of continuous aging were used with the permission of their creator, Mr. Anthony Cerniello. Images of a city skyline during different hours of the day were generated using the computer game “Watch Dogs 2” and were used in accordance with the publisher (Ubisoft) guidelines.

All subjects underwent a structured 4-day training routine (30). Each day subjects performed a series of four tasks. In Task 1, participants familiarized themselves with the mechanics of manipulating the age of the person and the time-of-day of the city backdrop. Subjects were presented with a series of randomly selected age-day combinations and were required to manipulate the adjacent interactive image’s age and/or time-of-day to match the assigned combination (Movie S2). Here, as in all subsequent behavioral tasks, morphing the interactive image was performed using a standard computer mouse, with horizontal motion controlling time-of-day (earlier: left, later: right) and vertical motion controlling the age (younger: down, older: up). To minimize the use of spatial strategy, the mouse cursor was made invisible during age-day manipulation.

In Task 2, to reinforce subjects’ ability to mentally simulate changes in age and time-of-day, they were, again, presented with randomly selected age-day combinations and were required to manipulate the adjacent image’s age and time of day to match the assigned combination. Specifically, in task 2 (Fig. S1B), subjects were able to adjust age-day only within a limited range around the initial age-day combination. To achieve their objective, once performing the (limited) age-day adjustment, subjects would repeat their adjustment automatically until the image had either matched the assigned combination or reached the limit of either age or time-of-day morphing (Movie S3). Subjects received feedback regarding their success in each trial.

In task 3, subjects established associations between the 9 age-day combinations and 9 event symbols. Subjects were randomly presented with each of the 9 embedded event symbols and were required to manipulate the adjacent interactive image’s age-day to match the associated age-day combination (Movie S4).

Combining multiple elements from all preceding tasks, in Task 4 subjects were presented with randomly selected event symbols and were required to manipulate the adjacent image’s age-day to match the embedded event’s age-day coordinates. Similarly to Task 2, subjects were limited in their ability to manipulate age-day, and consequently were instructed to adjust the age-day, such that if repeated automatically, the image would match the age-day coordinates associated with the selected event image (Movie S5). Subjects received feedback regarding their success in each trial. The stimulus-outcome tasks were presented using an in-house made Matlab GUI. Cursor location, response times and success rates were recorded. Importantly, to avoid priming the subjects towards spatial reasoning and strategy, instructions on the tasks were given while avoiding the use of any spatially-related language.

### fMRI-adapted task

Through training, subjects gained extensive experience in the regularities governing the age-day space as well as the embedded “event symbols”. Immediately following 4 days of training, subjects performed an fMRI-adapted task in the age-day space, analogous to tasks used for navigation in physical space (28, 60).

During the fMRI scan, subjects watched videos of an image continuously changing in age and time-of-day, according to predefined age:day ratios (30). Each video corresponded to one trajectory and consisted of several stages: (1) morphing, (2) imagination and (3) feedback (Fig. 2A,B, Movie S1).

During the morphing stage, subjects viewed a single age-day trajectory for 8 seconds. Next, during the imagination stage, subjects were presented with a blank screen and were instructed to further mentally simulate the image morphing with the same age:day ratio and at the same speed for an additional 4 seconds. Finally, during the feedback stage, subjects were presented with three outcome stimuli and were given 4 seconds to select the one associated with the trajectory’s outcome. Responses were given by pressing one of three keys on a controller. In total, subjects performed 132 such trials, equally divided into 3 runs, with 6 second intervals between trials.

### MRI data acquisition

For details regarding MRI data acquisition please refer to supplementary materials.

### MRI preprocessing

For details regarding MRI data preprocessing please refer to supplementary materials.

### Measures of motion estimation

Correction for head movement was performed at two levels: First, we applied the realignment algorithm of SPM12, which is specifically designed to account for head movement during fMRI scanning. This algorithm corrects for motion-related linear or angular displacement of scan images. Second, we included movement parameters for each scan volume (as calculated by the realignment algorithm) as regressors of no interest in every GLM that was carried out in order to calculate grid-cell-like representations. This approach corrects for movement-related signal artifacts (i.e., spin-history effects), so that other regressors in the GLM (e.g., like regressors testing for the 6-fold symmetric modulation) are not affected by movement-related changes of the BOLD signal.

### Behavioral data analysis

In the fMRI task, we computed the success rates (SR) as percentage of correct responses across all scanning sessions, and response time (RT) as the time from the start of the feedback stage to a button press. In addition, we examined the effect of timeline congruency on task performance. Trials were classified as “congruent” if trajectories had both age and time-of-day progress from young to old and from day to night, respectively, and “incongruent” otherwise. We compared congruent and incongruent trials using pairwise t-tests on SR and RT.

### Entorhinal cortex segmentation

Segmentation was conducted using the ITK-SNAP (http://itksnap.org; v.3.8.0) software (61). ITK-SNAP is a freely available open source DICOM-software tool for viewing and segmentation of MRI scans. Entorhinal cortices were automatically segmented according to the ASHS-PMC-T1 1.0.0 protocol (ITK-SNAP DSS).

### fMRI data analyses

With the intention of testing whether the fMRI signal in EC is characterized by hexagonal symmetry (as a proxy for grid cells activity) we conducted a three-part analysis using the Grid Code Analysis Toolbox (GridCAT) (62): (1) Using data from the first run (out of total three), we evaluated whole-brain hexagonal modulation, indicating a voxel’s tendency to correspond to a periodicity of 60°. Importantly, as this metric is indifferent to grid orientation, it was only used to create orthogonal ROIs that allowed us to test for grid-related activity in an unbiased fashion; (2) We defined ROIs as 6mm spheres centered at points of maximal hexagonal modulation within the personalized EC maps bilaterally; (3) Next, we split the remaining data (runs 2,3) in two, using the first half to estimate voxel-wise grid orientations in the EC ROIs and then testing these orientations on the other half of the data in order to quantify the magnitude of each subject’s grid-cell-like representations. As a control analysis, we repeated split-half procedure for two alternative symmetrical models (5- and 7-fold symmetry).

### Hexagonal modulation analysis

We modeled the fMRI time series of the first run using a general linear model (GLM, 64). The “morph”, “imagine”, and “feedback” stages were modeled using regressors for main effect and two parametrically modulated regressors (30): the sine and cosine of the direction of each trajectory in each trial, θ(t), with a periodicity of 60°, that is, sin(6θ(t)) and cos(6θ(t)), as well as six nuisance regressors to account for motion-related artifacts. Importantly, here, as in all further analyses, only parameter estimates of “imagine” condition in correctly replied trials were analyzed. We have analyzed only the “imagine” condition to minimize the direct influence of the stimuli’s visual properties on the grid-cell-like signal. Correct trials were selected as only these satisfy the fundamental association between the direction of trajection and the corresponding angle, that lies at the basis of the grid-cell-like analysis. We subsequently used the sine and cosine parameter estimates to generate maps of “firing amplitude” 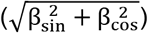, allowing us to quantify hexagonal modulation for each voxel.

### Defining ROIs

To increase the likelihood of correctly identifying regions with grid-like behavior we constructed 6-mm sphere ROIs, centered at the point of maximal amplitude, within the individually segmented EC maps of each subject (see Entorhinal cortex segmentation).

### Split-half analysis

First, we partitioned each of the two remaining fMRI runs (runs 2,3) into two halves, using the first half of each run as the estimation dataset and the second half as test dataset. We estimated voxel-wise grid orientations by fitting the estimation dataset to a first general linear model (GLM1). Similarly to the hexagonal modulation analysis, we used the direction of each trajectory (θ(t)), to construct a model that included two parametrically modulated regressors, sin(6θ(t)) and cos(6θ(t)) for the “morph”, “imagine”, and “feedback” stages of correct trials, while incorrect trials were modulated as non-grid events. We then calculated the mean grid orientation (ϕ) within each EC ROI, by averaging the “imagine” beta estimates (βsin and βcos) associated with the two parametric modulation regressors over all voxels in the left and right EC ROIs and assigning the resulting two values to: ϕ = arctan(βsin/βcos)/6. Then, the remaining half of the data (test dataset) was modeled in a second general linear model (GLM2), that included two regressors: “aligned” and “misaligned”. Aligned/misaligned status was determined by taking each event’s translation direction (θ(t)) and calculating its difference from the mean grid orientation (ϕ). An event was considered “aligned” if its event-angle lied within ±15 degrees of the mean grid orientation (or a 60-degree multiple of this value), and otherwise as “misaligned” (Fig. 2D). Finally, grid-cell-like representations were quantified by averaging “aligned” and “misaligned” parameter estimates within EC ROIs voxels. We performed a pairwise t-tests on the differences between the resulting mean aligned and misaligned parameter estimates, as well as on the differences between each of the parameter estimates and zero. As a control analysis, we repeated split-half procedure for two alternative symmetrical models (5- and 7-fold symmetry).

To determine whether coding of past-to-future congruency could modulate grid-cell-like representations we conducted a modified split-half analysis: First we divided right and left EC ROIs into voxels decoding congruency above (>0.5, “EC ∩ MVPA”) and (2) below (≤0.5, 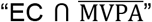) chance level using the results of the MVPA analysis (see “MVPA searchlight analysis”). We then estimated the mean grid orientation using the first half of the data from runs 2,3 separately for each of the ROIs. Subsequently, we used the second half of the data from runs 2,3 to construct four GLM2 models that estimated “aligned” and “misaligned” parameters, separately for each of the ROIs. The magnitude of grid-cell-like representations in each ROI was quantified by averaging the “aligned” and “misaligned” parameter estimates (of the “imagine” condition in correct trials only) across all ROI voxels. Finally, a pairwise t-test on the differences between the mean aligned and misaligned parameter estimates, was performed to determine whether concurrent congruency coding modulates the magnitude of grid-cell-like representations.

### MVPA searchlight analysis

Our secondary research question was whether timeline congruency could modulate grid grid-cell-like representations. For this purpose, we attempted to identify brain regions implicated in timeline congruency processing, using a searchlight-based MVPA method (39).

As a preliminary step, we performed a GLM fitting “congruent” and “incongruent” regressors to the time course, as well as six nuisance regressors to account for motion-related artifacts. Trials were deemed “congruent” if their trajectories progressed from both young to old and from morning to evening (i.e aligned with the natural progression of time), and “incongruent” otherwise (Fig. 4A,B).

Subsequently, we employed a support vector machine (SVM) classification algorithm implemented in the CoSMoMVPA toolbox for Matlab (64) on the parameter estimates of the congruency GLM analysis. Each voxel was taken as the center of a 3-voxel radius sphere. For each of those spheres, a 5-fold leave-one-out cross-validation was performed, resulting in an accuracy value per sphere. To assess the statistical significance of searchlight maps across participants, all maps were corrected for multiple comparisons without choosing an arbitrary uncorrected threshold using threshold-free cluster enhancement (TFCE) on the cluster level (65). A Monte-Carlo simulation permuting condition labels was used to estimate a null TFCE distribution. First, 100 null searchlight maps were generated for each participant by randomly permuting condition labels within each obtained searchlight classification. Next, 10,000 null TFCE maps were constructed by randomly sampling from these null data sets in order to estimate a null TFCE distribution (66), obtaining a group level Z-score map of the classifier results. Voxels surpassing a significance threshold (Z-score>1.645, P<0.05) were examined for overlap with the previously defined EC ROIs. The degree of overlap was calculated for both the central voxel for every sphere, and for the whole sphere.

## Acknowledgements

We thank Prof. Dori Derdikman (Technion – Israel Institute of Technology) and Dr. Michael Peer (University of Pennsylvania) for helpful discussions, and Prof. Tommy Kaplan (The Hebrew University) for invaluable support.

