## Supplementary Materials for "Evidence for grid-cell-related activity in the time domain"

### Results

#### Behavioral Training

In task 3, subjects freely morphed age and time-of-day to match the coordinates of embedded event symbols. Subjects' familiarization with the embedded outcome objects was revealed to have increased through training, as time-to-outcome gradually decreased throughout training and was significantly smaller for day four compared with day one (Fig. S1C, $F(3,68) = 4$, $P = 0.01$, one-way ANOVA with Scheffé post-hoc test).

In tasks 2 and 4, subjects were instructed to morph images into specific outcomes by determining an age:day ratio, thereby generating a continuous linear morph. Subjects' ability to simulate unseen outcomes increased as angle errors gradually decreased throughout training and were shown to be significantly different when comparing days 1 and 2,3,4 (Fig. S1A, $F(3,108) = 7.18$, $P < 0.0005$, one-way ANOVA with Scheffé post-hoc test) in task 2 and significantly different comparing days 1 and 4 (Fig. S1B, $F(3,108) = 2.85$, $P < 0.05$, one-way ANOVA with Scheffé post-hoc test) in task 4.

To evaluate how growing familiarity with the age-day space had manifested in task 3, we estimated the time it took subjects to navigate to the symbol images (time-to-outcome) and compared average time-to-outcome values between the different sessions across all subjects using one-way ANOVA with Scheffé post-hoc test.

Combining multiple elements from all preceding tasks, in task 4 subjects were presented with randomly selected event symbols and were required to manipulate the adjacent image's age-day to match the embedded event's age-day coordinates. Similarly to task 2, subjects were limited in their ability to manipulate age-day, and consequently were instructed to adjust the age-day, such that if repeated automatically, the image would match the age-day coordinates associated with the selected event image. (Movie S5). Subjects received feedback regarding their success in each trial. The stimulus-outcome tasks were presented using a custom-designed Matlab GUI. Cursor location, response times and success rates were recorded. Importantly, in-order to avoid priming the subjects towards spatial reasoning and strategy, instructions on the tasks were given while avoiding the use of any spatially-related language.

In tasks 2 and 4, to estimate performance we computed angle errors, defined as the angle between the ideal trajectory angle and the angle formed by the adjusted age:day ratio. Angle errors were compared between the different sessions and across all subjects using one-way ANOVA with Scheffé post-hoc test.

**MRI data acquisition**

Subjects were scanned in a 3T Siemens Skyra MRI (Siemens, Erlangen, Germany) at the Edmund and Lily Safra Center (ELSC) neuroimaging unit. Blood oxygenation level-dependent (BOLD) contrast was obtained with a gradient-echo, echo-planar imaging (EPI) sequence [repetition time (TR), 1000 ms; echo time (TE), 35 ms; multiband acceleration factor of 4; flip angle, 62°; field of view, 192 mm; matrix size, 96 × 96; functional voxel size, 2 × 2 × 2 mm; 44 slices, descending acquisition order; slice thickness, 2mm; gap, 0.2mm]. In-order to minimize signal loss in the orbitofrontal cortex region, slice angle was set to 30° relative to the anterior-posterior commissure (ACPC) line (2). In addition, T1-weighted high resolution (1 × 1 × 1 mm, 160 slices) anatomical images were acquired for each subject using the MPRAGE protocol [TR, 2300 ms; TE, 2.98 ms; flip angle, 9°; field of view, 256 mm]. Stimuli presentation and subject's button presses were registered and time-locked to the fMRI data.

**MRI preprocessing**

For details regarding MRI data preprocessing please refer to supplementary materials.

fMRI data were analyzed using the SPM 12 software package, version 7219 (3) and in-house Matlab (Mathworks) scripts (available at neuropsychiatrylab.com). Preprocessing of functional scans included slice-time correction (sinc interpolation), 3D motion correction by realignment to the first run image (2nd degree B-Spline interpolation), exclusion of runs with maximal motion above a single voxel size (3 mm) in any direction, smoothing (full width at half maximum (FWHM) = 4 mm), and co-registration to the anatomical T1 images. Anatomical brain images were corrected for signal inhomogeneity and skull-stripped. All images were subsequently normalized to Montreal Neurological Institute (MNI) space (3 × 3 × 3 mm functional resolution, 4th degree B-Spline interpolation).

**Hexagonal modulation analysis**

As a complementary analysis to our ROI-driven approach, we conducted a whole-brain analysis to identify voxels showing significant hexagonal modulation in a hypothesis-free manner. For this purpose, we conducted a group GLM of $\sin(6\theta)$ estimators and included task success rate as covariate. Whole brain group analysis showed significant hexagonal modulation in the temporal pole bilaterally, right hippocampus and left EC (Fig. S2).

**Split-half analysis**

To exclude any potential bias caused by imbalanced sampling due to analyzing correct trials only, we have analyzed the distribution of aligned and misaligned correct trials. We found no statistically significant difference between the number of trials in the aligned and misaligned conditions (paired t-test: $t(27) = 0.68$, $p > 0.05$).

Supplementary movies can be found at:

https://drive.google.com/drive/folders/10U1I4Alv4BRBZ9k2uji5G3d4HmeQc273?usp=sharing

**Figures**

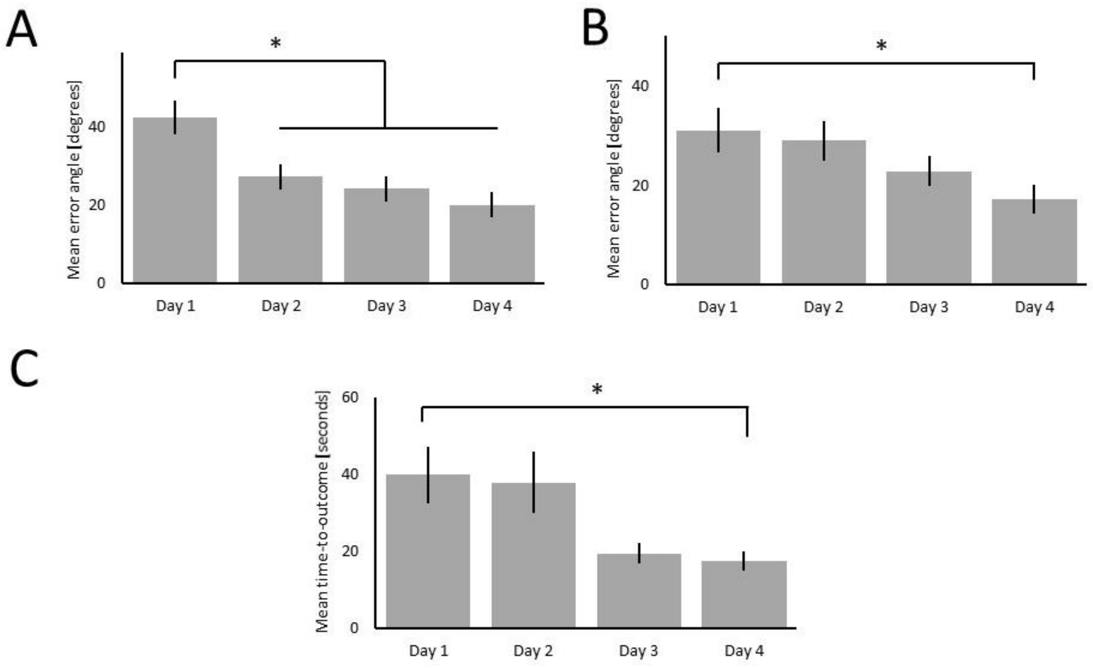


**Figure S1. Behavioral performance during training.** (A) Performance in training task 2. Barplots show the mean error angle, defined as the angle between the optimal trajectory to an age-day combination and the trajectory generated by the subject-adjusted age:day ratio, to gradually decrease throughout training. Mean error angle is shown to significantly differ between day one and days two, three and four ($P<0.05$, ANOVA and Scheffé post hoc). (B) Performance in training task 4. Barplots show the mean error angle, defined as the angle between the optimal trajectory to an event symbol and the trajectory formed by the subject-adjusted age:day ratio, to gradually decrease throughout training. In task 4, Mean error angle is shown to significantly differ between day one and day four ($P<0.05$, ANOVA and Scheffé post hoc). (C) Estimating familiarization with the embedded outcomes. In training task 3, we estimated the time it took subjects to “navigate” to an event symbol (time-to-outcome) and compared average time-to-outcome values between the different sessions. Result show time-to-outcome gradually decreased throughout training and was significantly smaller for day four compared with day one ($P<0.05$, ANOVA and Scheffé post hoc).

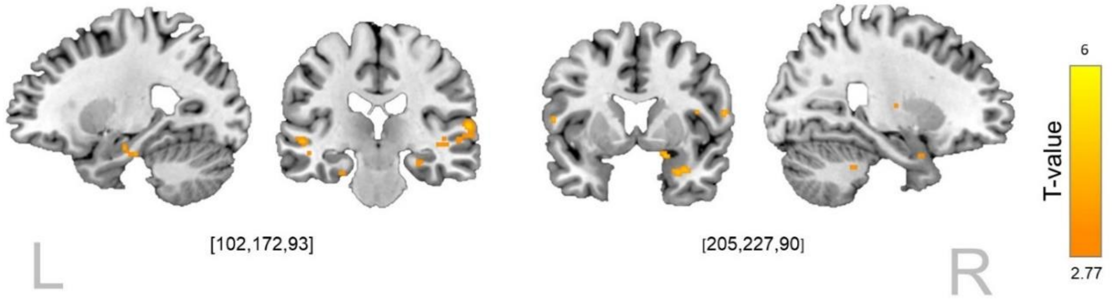


**Figure S2. Identifying hexagonally symmetric signals across the whole brain.** Whole-brain parametric analysis of first run data showed significant estimators of the sin(6θ) and task success rate covariate were found at the right Putamen, Temporal pole, Hippocampus and EC bilaterally. ($P<0.005$, uncorrected, 5 voxel cluster size thresholding)
